# Accurate Reconstruction of Adaptive Immune Receptor Repertoires from FACS-partitioned B and T Cell Subsets

**DOI:** 10.64898/2026.09.13.746584

**Authors:** Hahn Zhao, Alexandra Morgan, Michie Yasuda, Hamid Mirebrahim, Ulrich Schlecht, Sylvie McNamara, Rena Adachi, Sandra Siemann, Florian Rubelt, Dinesh Kumar, Sowmi Utiramerur, Ramy Arnaout, Hosseinali Asgharian

## Abstract

Adaptive immune-receptor repertoire sequencing (AIRRseq) is crucial for understanding immune system diversity and its relationship to disease dynamics. Partitioning of total B and T cells into their major subsets with distinct immunological functions—IgM+ vs. class-switched B cells (IgG+ > IgA+) and CD4+ vs. CD8+ T cells, respectively—allows for AIRRseq-based analysis of the unique contributions of each compartment to the overall immune response, a major advantage over traditional bulk sequencing workflows. However, data from these subsets is not directly comparable with the vast majority of publicly available AIRRseq data, which comes from unfractionated B and T cells, an important incompatibility. Here we investigate computational methods for reconstructing complete AIRRseq repertoires from partitioned B and T cell subsets in diverse individuals. Peripheral blood mononuclear cells (PBMCs) were partitioned via positive selection of IgM+ B-cell subsets and CD4+ T cells using immunomagnetic beads; genomic DNA was then extracted and B- and T-cell receptors were sequenced. Four reconstruction methods are introduced and evaluated for concordance with matching unpartitioned repertoires to assess preservation of key repertoire characteristics. Results show that these methods enable accurate estimates of overall immune-repertoire diversity from B- and T-cell subsets in a way that simply pooling the sequence data from sub-repertoires cannot.

## Background

The adaptive immune system recognizes and reacts to a wide array of antigens due to the diversity of the B- and T-cell receptor (BCR and TCR) repertoire, commonly referred to as the immunome. Repertoires exhibit modest overlap between individuals at the sequence level (Robins et al. 2010). At the same time, higher-level patterns across individuals often correlate with specific antigenic exposures and diseases (Mark et al. 2022). Identifying these shared signatures requires evaluating extensive sequence datasets from multiple studies. Consequently, standardization methods are needed to construct consistent qualitative and quantitative immune repertoire models that can be seamlessly integrated across diverse experimental platforms and analytical pipelines.

B and T cells are functionally heterogeneous, with subsets of these cells performing distinct tasks. New B cells express antibodies of the IgM class or isotype, a pentameric and generally low-specificity form, switching to high-specificity IgG (or other classes) after antigen exposure (Carsetti et al. 2022). CD4+ helper T cells differ from CD8+ cytotoxic T cells, with the former playing more regulatory roles (Laidlaw et al. 2016). Of note, the relative sizes of these subsets carry clinical information and can vary by condition and age. For example, the CD4:CD8 ratio is a useful marker of immunosuppression, and is an important clinical measurement in the setting of HIV infection (Miller et al. 2025). Additionally, there has been much interest in diversity indices such as species richness, Shannon entropy, and Simpson’s index, especially as unified under Hill’s and more recently Leinster, Cobbold, and Reeve’s diversity-number frameworks (Hill 1973, Leinster & Cobbold 2012, Reeve et al. 2016). This is because these indices are known to differ under different physiological, pathophysiological, and treatment states, for example in aging and during cancer treatment (Akbari et al. 2025). Identification of relative abundance, diversity and individual B and T cell clones involved in these specialized functions requires separate sequencing of each functional compartment.

Bulk quantitative sequencing remains the most common, largest-scale, and most cost-effective approach to AIRRseq, using either DNA or mRNA as starting input material (Seo and Choi 2025). mRNA is more plentiful but biases the results toward transcriptionally active cells, while starting with DNA facilitates enumeration of all cells in an unbiased manner (Mazzotti et al. 2022). This is critical especially for calculating quantitative representations of the immune systems such as the various metrics of diversity and cross-sample similarity or distance. For this reason, we focus on DNA-based immune profiling in this study. For either input, the starting sample is most often whole blood obtained via venipuncture (peripheral blood). Density-gradient centrifugation is regularly used to deplete red blood cells and neutrophils from whole blood. The resulting subset, known as peripheral blood mononuclear cells (PBMCs), usually comprises 50-80% lymphocytes (B and T cells) which are the target cells for AIRRseq assays (Kleiveland 2015). PBMCs can then be partitioned into B- and T-cell subsets or sequenced without partitioning. When partitioning is performed, affinity purification columns are more cost-effective than flow cytometry and much more cost-effective than single-cell sequencing, making columns better suited for large-scale studies. Once the desired subset is obtained, cells are lysed, the DNA is bulk-purified, the VDJ-recombined BCR or TCR DNA is barcoded for quantitation (through limited PCR amplification), and a next-generation sequencing (NGS) library is prepared, normalized, and sequenced using an established AIRRseq workflow.

The value of sequencing specific, surface-marker-defined lymphocyte subsets is increasingly evident in clinical oncology and infectious disease research. For instance, targeted TCR sequencing in subjects with non-small cell lung cancer revealed that high pre-treatment diversity within peripheral PD-1+ CD8+ T cells, a potentially tumor-reactive subset, serves as a strong, non-invasive predictor of improved progression-free survival and favorable response to immune checkpoint blockade (Gelibter et al. 2024). In another study, PD-1 expression was used to identify subject-specific, tumor-reactive CD8+ T cell repertoires (Gros et al. 2014). In this study, deep sequencing exclusively on isolated CD8+ PD-1+ tumor-infiltrating lymphocytes (TILs) revealed that the most highly-expanded clonotypes within this exhausted subset directly recognized autologous tumor antigens. Beyond oncology, this partitioning principle is equally valuable for autoimmune conditions. For example, isolating peripheral CD4+ T cells in people with type 1 diabetes revealed distinct, disease-associated CDR3 overlap patterns that are otherwise averaged out by more abundant subsets (Gomez-Tourino et al. 2017). In all these cases, biological signatures would likely be obscured in an unpartitioned bulk analysis.

To date, most public AIRRseq data is for “overall” repertoires, i.e. from non-partitioned cells (Olsen et al. 2022; Corrie et al. 2018). This data includes some very large studies that make useful comparands. Given that bulk (as opposed to single-cell) subset studies are increasingly common, the question arises of how to compare data from repertoires that are sequenced as subsets to data from overall repertoires. Even though bulk sequencing is usually quantitative, meaning each sequenced cell is barcode-enumerated, as a rule the overall repertoire cannot dependably be recovered simply by pooling the data from the subsets. This is because of normalization during next-generation sequencing (NGS) library preparation, during which step information about the relative sizes of the subsets is lost. Additionally, because diversity indices can be sensitive to sample size—i.e., the number of cells sequenced—and because of details of their mathematical formulations, diversity indices for the overall repertoire are usually not just the sums or averages of the diversities of the subsets. However, it is possible to estimate properties of an overall repertoire from AIRRseq data on its subsets if appropriate steps are taken during the analysis. This work presents and evaluates computational/bioinformatic methods to this end.

## Materials & Methods

### Subject Selection

All work was approved by the Institutional Review Board of the Beth Israel Deaconess Medical Center (protocol no. 2021P000899). EDTA-anticoagulated whole blood was collected from 30 subjects: 11 diagnosed with cancer (pancreas, breast, prostate, colorectal, or liver), nine with bacterial blood infections (with the clinically important pathogens *Enterococcus faecalis, Enterococcus faecium, Pseudomonas aeruginosa, Klebsiella pneumoniae*, or *Escherichia coli*), and 10 healthy controls.

### Isolation and Separation of T-Cell Subsets

For each subject, PBMCs were isolated from 3 mL of blood within 10 hours of blood collection. Isolation was performed using Ficoll-Paque PLUS density gradient media (Cytiva, Little Chalfont, United Kingdom) and SepMate-15 tubes (Stemcell Technologies, Vancouver, Canada). The isolated PBMCs were then resuspended in CELLBANKER 2 (AMSBIO, Oxfordshire, UK) and stored frozen at −80°C until use.

Upon thawing and removal of the cell-freezing medium, cells were resuspended in cold autoMACS Running Buffer (Miltenyi Biotec, Bergisch Gladbach, Germany). One third of this suspension was used as a source for DNA from the “total cell” fraction (TL), which contained all of the B and T cell subsets in this work: a sample of the unfractionated/overall repertoire. A small aliquot was set aside to perform a cell differential count (henceforth “differential”) via fluorescence-activated cell sorting (FACS), providing the ground-truth count of B and T cells and their subsets. The remaining PBMCs were divided into two fractions using immunomagnetic human Anti-IgM MicroBeads and CD4 MicroBeads on Multi-24 Column Blocks using the MultiMACS Cell24 Separator Plus (Miltenyi Biotec, Bergisch Gladbach, Germany). These are referred to as “labelled cells” (LC), containing IgM+ B cells and CD4+ T cells; and “flowthrough” (FT), containing non-IgM+ B cells (mostly IgG+ but also IgA+ and rare IgD+ and IgE+) and CD8+ T cells **(Figure 1A)**. The entire LC fraction was used for DNA extraction, while a small portion of FT was removed for differential by FACS before DNA extraction. DNA was extracted and purified using the Mini spin kit for ultrafast DNA isolation from blood (MACHEREY-NAGEL, Düren, Germany). Elution of DNA was performed using 10 mM Tris-HCl buffer (pH 8) instead of the kit’s elution buffer (EB).

**Figure 1.**
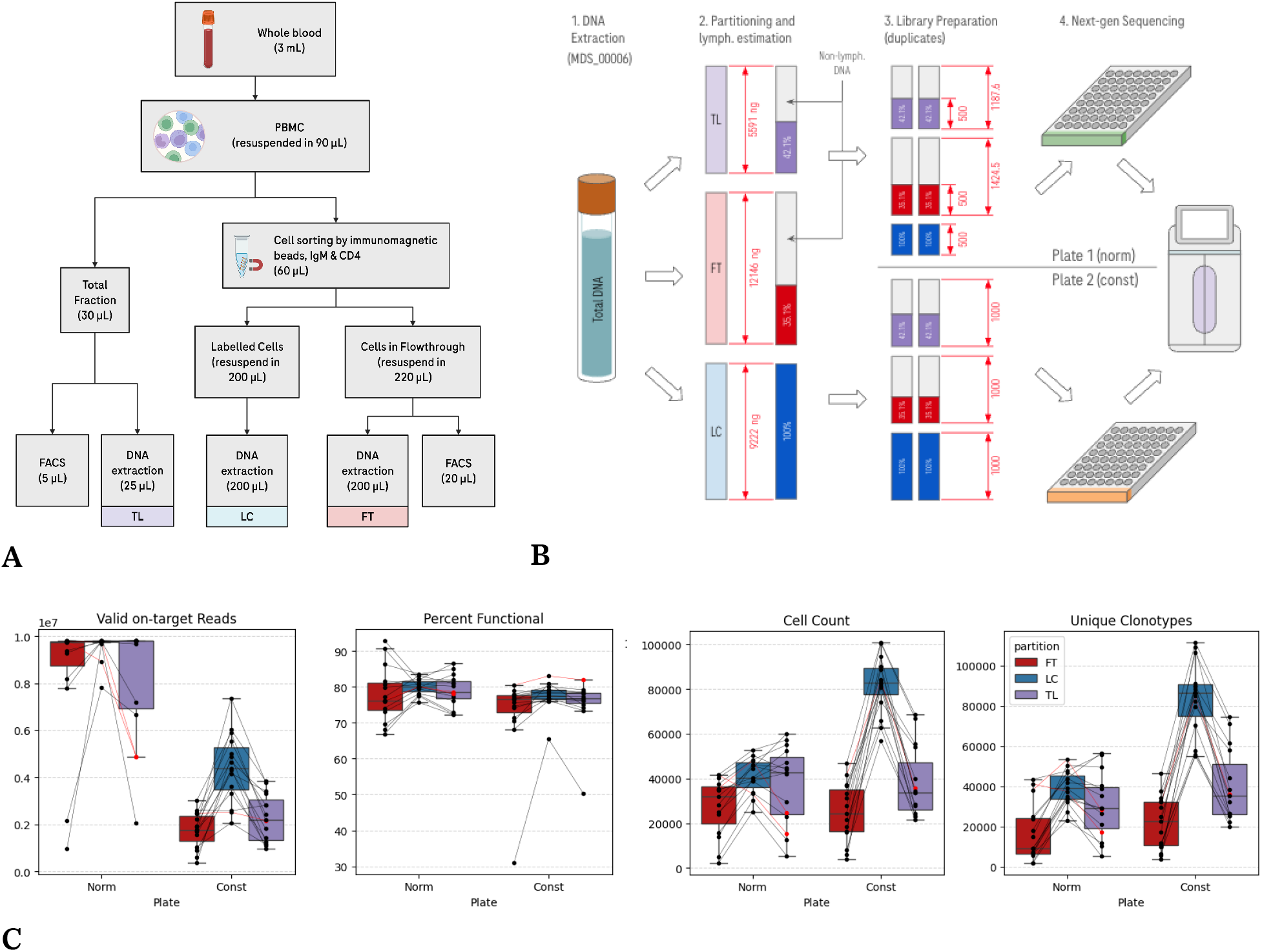
**(A)** Schematic diagram outlining the workflow for PBMC isolation, cell sorting, and partitioning into total, labeled, and flowthrough fractions for downstream analysis. **(B)** Overview of plate design and lymphocyte partitioning: DNA is partitioned into TL, FT, and LC fractions for each of the two plate designs Norm and Const), followed by library preparation and sequencing. Red annotations mark anticipated nanograms of lymphocyte DNA at each step. Two reconstruction methods (“match” and “equal”) were applied to aggregated repertoires, highlighting differences in the allocation of LC and FT contributions. **(C)** Initial quality control metrics for partitioned repertoires across each of the two experimental plates (Norm and Const). Partitions that did not have sufficient DNA input for a full set of replicates are marked in red. Normalizing the total input DNA based on lymphocyte fraction in Norm results in more equally represented cell recovery across partitions compared to the un-normalized setup in Const.

### Measuring lymphocyte content in FACS-sorted cells

To quantify cell fractions in FACS-sorted cells, genomic DNA was extracted from 0.83 mL of blood in the TL partition, 1.82 mL in FT, and 2 mL in LC for each subject. UV/vis spectrophotometry was used to estimate total DNA and lymphocyte DNA concentrations from the post-elution buffer on a 20.5 µL sample. The proportion of CD4+ or IgM+ target DNA in the subject (w_LC_) was estimated as the lymphocyte DNA concentration in LC expressed as a fraction of total lymphocyte concentration ([DNA FT] + [DNA LC]). The full FACS data is presented in **Table A1**.

### Sequencing plate design

TL, LC, and FT fractions from the 30 subjects were divided across two 96-well plates **(Figure 1B)**. To test whether adjusting input mass to lymphocyte content improves library fidelity over fixed-input workflows, samples were divided across the two plates representing lymphocyte-normalized (‘Norm’) and constant total DNA (‘Const’) protocols. On plate Norm, adjusted to 500 ng of lymphocyte DNA. On plate Const, the total input DNA was set to 1,000 ng, which includes the DNA of any non-lymphocyte white blood cells present in the fraction. If the 1,000-ng target was not met, all available DNA was used, with a minimum threshold of 300 ng. Efforts were made to minimize the difference in fraction of lymphocyte DNA in the FT partition across the two plates. For each subject, BCR and TCR libraries were prepared as biological duplicates for each of the three fractions. Three of the 30 subjects (two Norm, one Const) had sufficient extracted DNA for only a single TL replicate.

### DNA sequencing and data pre-processing

Rearranged BCR and TCR genomic DNA (gDNA) was captured using a multiplexed immune repertoire primer extension and target enrichment kit developed by Roche (Donahue et al. 2019). This kit produces libraries that include immunoglobulin heavy chains for BCR (IGH) and beta and delta chains for TCR (TRB and TRD). Resulting libraries were sequenced using 2×150-base paired-chain reads on Illumina’s NovaSeq platform at Signature Diagnostics/Roche in Potsdam, Germany. A total of 190 libraries, including 7 blanks and 6 PBMC controls (StemCell Technologies Cat. No. 70025) generated a total of 4.7B reads, with approximately 25.6M reads per library in clinical samples.

V/J-gene annotations and CDR3 sequences were extracted from raw sequencing data using the Daedalus bioinformatics pipeline developed by Roche. **(Figure S1)**. In brief, raw FASTQ sequences were assessed for UMI pattern validity and down-sampled to a maximum of 50 million valid reads per library. Read mate pairs were trimmed of primer templates and aligned to annotated IG and TR references from the IMGT database (Lefranc et al. 2009). Gene-aligned reads were filtered to a threshold quality score of Q30, and 5 million on-target reads were subsampled and clustered based on CDR3 content and UMI sequence into UMI families.

Classification of productive UMI families was applied under the following conditions: (1) non-chimeric V-gene and J-gene annotations (i.e., families with V-gene and J-gene aligned to different loci); (2) absence of premature stop codons, ambiguous bases, and off-frame alignments; and (3) a CDR3 length of 10–24 amino acid residues, consistent with a study by Qian et al. that accounted for 97.0% of all families (Qian et al. 2024).

## Results

### Estimations of LC/FT fractions with pairwise overlap

To estimate the true proportions of labelled cells (LC) and flow-through (FT) fractions, we calculated the Bray-Curtis similarity index using the intersection (e.g. number of cells with identical clonotypes) between partitions as a fraction of the combined cell count in both groups **(Equation 1)**. The measurement uses the number of cells that share the same clonal rearrangement between samples A and B, where *n* is the number of unique clonotypes in both samples, and *f* is the frequency of clone *i*. Overlap between each partition and the unfractionated TL were used to determine the relative contribution of each partition, which was then used to predict the LC fraction (w_LC_). To evaluate precision and accuracy of estimations, we determined the concordance correlation coefficient (CCC) in **Equation 2** to measure the consistency of estimated *w*_LC_ using their correlation and deviation from the diagonal line of perfect agreement (Akoglu 2018).

We determined that in Norm where all partitions have equal target inputs, the highest concordance (CCC = 0.79) was achieved using gene usage counts as the unit of measurement. In order to mitigate the larger difference in target lymphocyte inputs in Const, raw CDR3 counts were normalized based on relative abundance, resulting in a more modest CCC of 0.67 **(Figure 3A)**.

**Figure 2.**
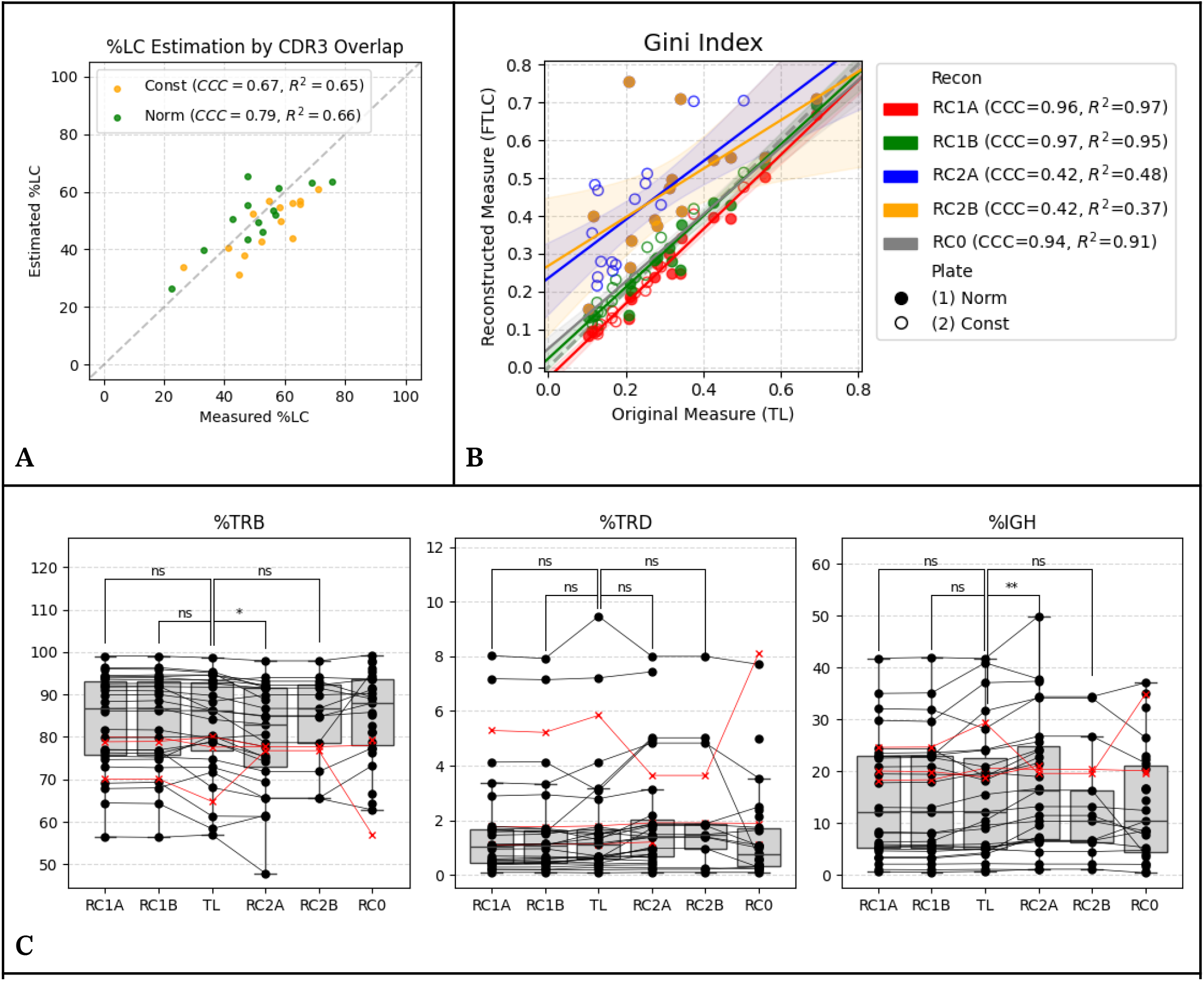
**(A)** Test of concordance for estimation of w_LC_ using relative weighted overlap between each partition and TL. CDR3 clonotype abundance was a better estimator in Const where partitions were set to a constant total DNA input (CCC = 0.67), whereas V/J-gene usage counts showed improved prediction when partitions were balanced by target input DNA in Norm (CCC = 0.79). **(B)** Gini dominance index of RC1 shows higher concordance than RC2 against TL. There was no notable improvement in the method RC1A over RC1B, despite the additional layer of subsampling to match the total TL input amount. **(C)** There was no significant difference in lymphocyte subtype ratios between either of the RC1 reconstructions and TL (paired t-test p ≥ 0.05), however, the RC2 method retains more rare clones, particularly those belonging to the less frequent TRD and IGH subtypes. This results in an overestimation of %TRD and %IGH compared to the TL repertoire.

**Figure 3.**
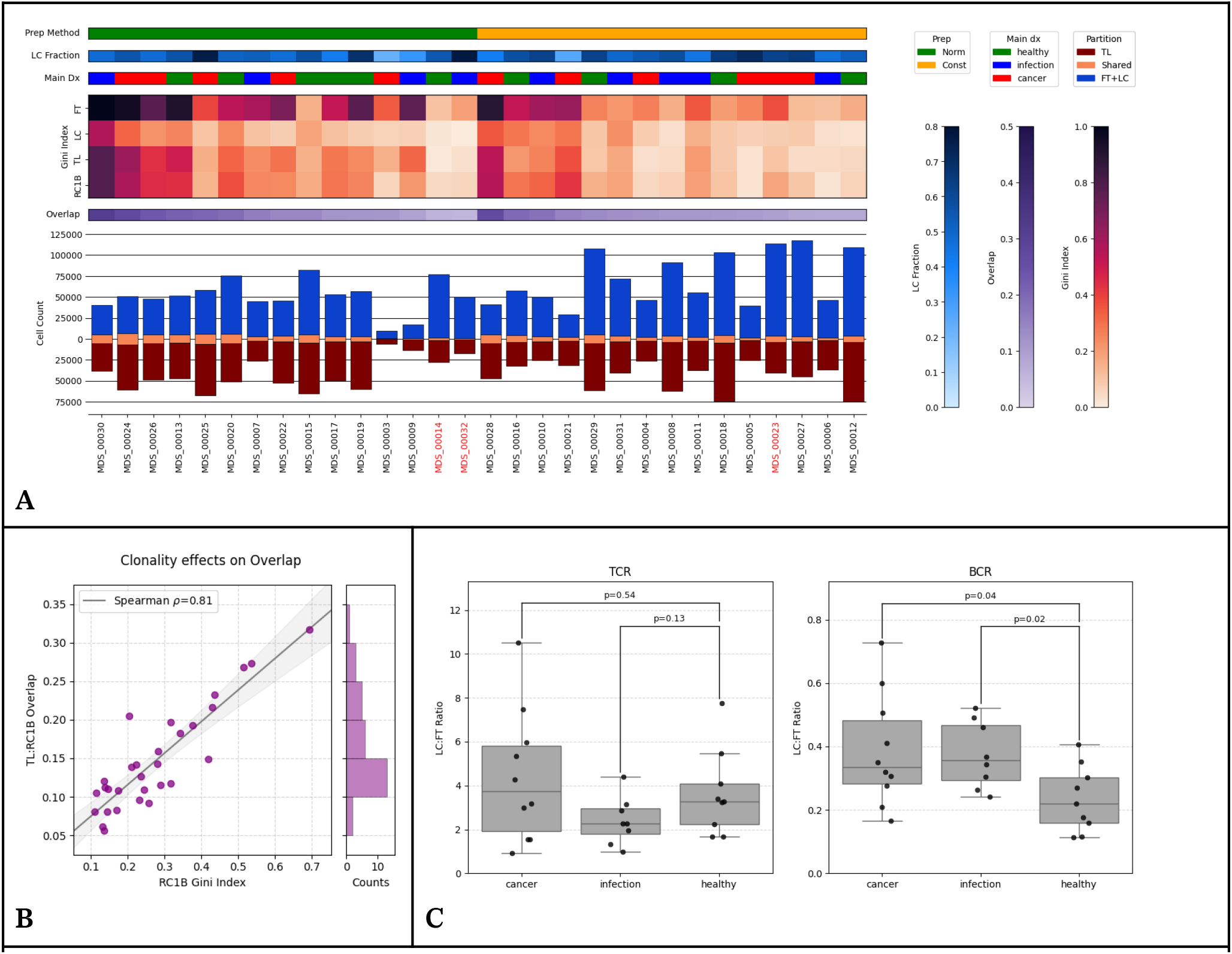
**(A)** Comparison of sample metrics between original and reconstructed repertoires. The divergent bar chart shows the number of cells in the assigned partition (RC1B in blue, TL in brown), and the number of shared cells in orange. **(B)** Sample overlap between TL and RC1B moderately correlates with repertoire clonality (ρ=0.81), due to the larger presence of abundant clones which are more likely to be captured across different replicates. **(C)** Comparison of reconstructions illustrating the ratio of TCRs (left) derived from the LC subset that underwent CD4+ selection relative to the unselected FT subset. Although not statistically significant, the infection group exhibits reduced LC ratios, suggesting a shift in immune dynamics favoring CD8+ cytotoxic T-cell activation. On the right, significantly lower values B-cell ratios from IgM selection in healthy subjects compared to cancer and infection subjects (ind. t-test p < 0.05) reflects a reduction in IgM-producing B-cells which are involved primarily in early immune responses.

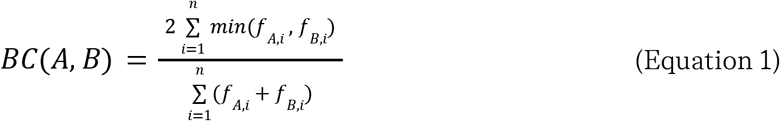

*The Bray-curtis similarity index is measured as the fraction of cells that share the same clonal rearrangement between samples A and B, where n is the number of unique clonotypes in both samples, and f is the frequency of clone i*.

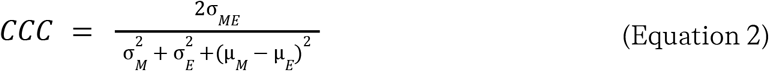

*The concordance correlation coefficient (CCC) measures the agreement between two continuous variables, in this case FACS-Measured (M) and Bioinformatically-Observed (O) evaluating the extent to which they deviate from the line of perfect concordance (M=E), thereby simultaneously assessing both bias and precision of the measurements*

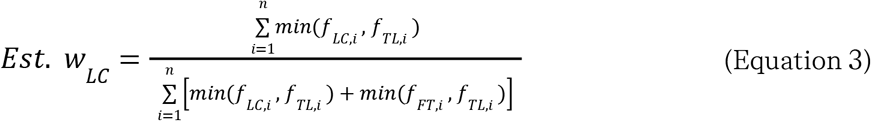

*Weighted clonal overlap between two samples can be defined as the number of cells in each sample that contribute to the union, expressed as a fraction of the total cell count across both samples*.

### Reconstructing repertoires from LC/FT fractions

#### Comparison of cell-level subsampling (RC1) and assigning clone weights (RC2)

Two high-level approaches to repertoire reconstruction were evaluated: sampling the cells of LC and FT repertoires at a ratio that matches *w*_LC_ and *w*_FT_ (RC1), and converting LC and FT repertoire cell counts into adjusted weights by using *w*_LC_ and *w*_FT_ as scaling factors (RC2). As a reference (RC0), LC and FT partitions were combined without any form of sampling.

For RC1, random subsampling was first applied to construct a joint repertoire that is normalized to match the lymphocyte amount in TL (RC1A, Equation 4). Second, a method agnostic of lymphocyte content in TL (RC1B, Equation 5) was attempted with the aim to retain as many clones as possible while maintaining the LC:FT ratio.

In contrast, for RC2, all observed clonotypes were retained while their cell counts were converted to non-integer weighted values. (i.e. *f*_*i*, RC_ = *f*_*i*, orig_ × *w*, where *f* is the frequency of clone *i*, and *w* is the weight factor representing the fraction of input DNA amount of a partition). All six replicates per subject were utilized in the RC2A reconstruction, whereas RC2B selects one FT and one LC for comparison with two replicates of TL. RC2B was only valid for subjects sequenced in plate-Norm, where libraries were balanced at an input amount of 500 ng lymphocyte content per replicate. Three subjects that did not have a full set of TL replicates (MDS_00014, MDS_00023, MDS_00032) were omitted from the analysis.

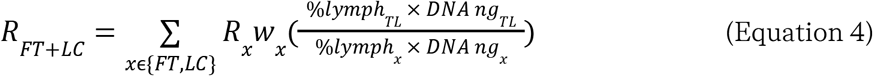

*R*_*FT+LC*_ *represents the reconstructed repertoire cell count as the combined size of its R*_*FT*_ *and R*_*LC*_ *components. In construct RC1A, families from each partition were subsampled to ensure proportional representation in the reconstruction, normalized based on the total amount of lymphocyte content in TL*.

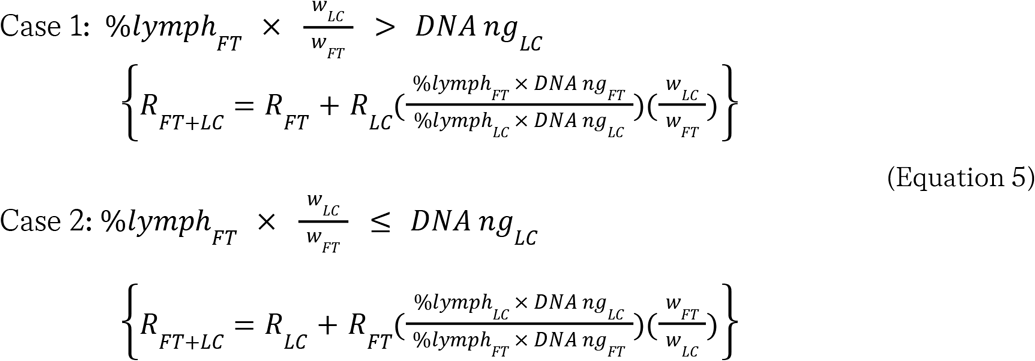

*In construct RC1B, the partition with DNA input constraints (e*.*g. FT in the equation above, LC below) is prioritized to ensure full representation retaining all cell counts, whereas the opposite partition is downsampled such that the joint FT+LC construct contains FT and LC repertoires appropriately sized to match w*_*FT*_ *and w*_*LC*_ *ratios*.

### Quality control and functional classification

We sequenced BCR and TCR in labeled cells (IgM+ B cells and CD4+ T cells), flowthrough (IgG+ B cells and CD8+ T cells), and the combined/total cell fractions—LC, FT, and TC, respectively. Overall, 77.4% of all unique receptor sequences were labeled as functional, consistent with expectations. The distribution of CDR3 lengths in productive and non-productive families is shown in **Figure S2**. The most common reasons for non-productive CDR3s were frameshift mutations resulting in a misaligned reading frame (16.5%) and premature stop codons (5%). Of the recovered lymphocytes in the unfractionated TL samples, the average composition was 81.8% ± 11.7% αβ T cells, 1.9% ± 2.2% γδ T cells, and 16.3% ± 12.0% B cells. FT and LC repertoires showed significantly different characteristics in cell type ratios (e.g. %TRB, %TRD, %IGH); specifically, the LC fraction had a higher proportion of TRB (93.8% ± 5.3%) compared to FT (62.7% ± 20.4%) **(Figure S3)**.

#### LC/FT-specific V-gene and J-gene usage patterns

To assess the impact of partitioning on immune repertoire gene usage, cell count frequencies for all possible V/J rearrangements were tabulated for each subject and normalized to counts per million **(Figure S6)**. To evaluate gene usage patterns across different health states, mean usage frequencies were further stratified into three groups: (1) healthy, (2) infection, and (3) cancer **(Figure S7)**. Both the infection and cancer groups showed significantly more differentially abundant V/J rearrangements in the LC partition, suggesting a likely shift in clonal expansion of select CD4+ T cell subpopulations in response to encountering disease-associated antigens.

### Comparing four repertoire reconstruction methods

We evaluated four methods for reconstructing overall repertoires from LC and FT fractions: RC1A, RC1B, RC2A and RC2B. We found that RC1 methods maintain TRB, TRD, and IGH percentages (paired t-test p>0.05), using the percentages in TL as ground truth. In contrast, RC2 methods, which are designed to retain less abundant clones, had the tendency to overestimate rare TRD chains **(Figure 2C)**.

#### Assessment of key diversity metrics

Hill numbers, also called D numbers and written *D*_*q*_, are a series of diversity indices that quantify the effective number of clones in a community, accounting for the frequency distribution (Hill 1973). The parameter *q* controls the extent to which smaller clones are downweighted proportional to their size, with q typically ranging from 0 to infinity. Most common diversity measures map to a certain value of *q*. For example, *D*_0_ is the species richness, describing the number of unique clones without regard to clone frequency. *D*_1_ corresponds to Shannon entropy (some downweighting of smaller clones), *D*_2_ corresponds to Simpson’s index (more downweighting), and *D*_∞_ corresponds to the Berger-Parker index, which reflects the relative abundance of only the most dominant clone (maximum downweighting).

Clonal diversity was measured through Hill numbers ranging from *D*_0_ to *D*_3_ and *D*_∞_ for each subject. Reconstruction methods RC1B and RC2A, which favor retention of rare clonotypes, led to greater diversity at lower Hill numbers **(Figure S4)**. D-number diversities for reconstructed subsets agree with TL for *q*≥2, suggesting that reconstruction captures larger clonotypes better than rarer ones.

The cumulative fraction of unique clones is plotted against the cumulative cell count in **Figure S5.2**. Known as the Lorenz curve, this curve measures inequality in the clone size distribution as deviation from the diagonal; the diagonal indicates a perfectly uniform repertoire, and the area between the curve and the diagonal as a fraction of the area beneath the diagonal is the Gini index, which is closely related to *D*_2_. As expected, the Gini index of RC1A and RC1B repertoires aligned strongly with TL measurements, with CCC scores of 0.96 and 0.97, respectively **(Figure 2B)**. Together, these findings support RC1B as the best of the four methods.

### Detailed comparison of reconstructions with TL

In 28 of the 30 subjects, the LC partition—consisting of selected CD4+ helper T-cells and IgM+ B cells measured lower in repertoire clonality compared to FT **(Figure 2C)**. Furthermore, we also observe differences in lymphocyte subtype ratios across the two partitions **(Figure S3)**. We sought to adjust the imbalance in repertoire diversity, and variations in lymphocyte subtype ratios across partitions (e.g. %TRB, %TRD, %IGH) during reconstruction through weighted sampling. We observed that repertoire clonality is directly proportional to sample overlap. This phenomenon occurs because as replicates are sequenced deeper, the dominant clones approach their steady-state abundance, whereas more rare clones are discovered with added depth **(Figure 3B)**.

The Bray-Curtis index for sample overlap measures the fraction of cells with shared clonotypes across two samples. We calculated the Bray-Curtis index between RC1B-reconstructed repertoires and unfractionated TL. Exact matching of clonal rearrangements is expected to result in overlap estimates near zero between subjects, as immune repertoires are highly disjoint. The mean index between TL and RC1B repertoires was measured at 0.130 ± 0.051 and 0.162 ± 0.076 for plates Const and Norm, respectively. For context, the mean overlap index was measured at 0.182 ± 0.145 in Const, and 0.353 ± 0.169 in Norm between technical TL replicates. Our findings suggest that adjusting DNA input amount based on lymphocyte abundance in a sample (i.e., using plate Norm) improves repertoire reconstruction fidelity, as evidenced by the higher overlap between the reconstructed repertoire and the ground truth.

### Exploring surface marker ratios in size-matched repertoires

Reconstructed repertoires offer a window into subset-specific immune dynamics by inferring immune marker ratios that would have been otherwise obscured through bulk sequencing alone. Expression of CD4+ and CD8+ markers can differ significantly in alpha-beta compared to gamma-delta T-cell subsets. The CD4:CD8 T-cell ratio typically ranges between 1.2 and 2.5 among healthy individuals, and is an important marker that can be traced to the level of stress encountered by the immune system. (Miller et al. 2025) Through determining the relative contributions of FT and LC in RC1B-reconstructed repertoires, we found lower CD4:CD8 T-cell ratios in infection groups, which could be indicative of CD8+ cytotoxic T-cell expansion in response to acute viral infections **(Figure 3C)**.

Similarly, IgM-to-non-IgM ratios were quantified in RC1B-reconstructed repertoires to assess isotype dynamics. B-cell repertoires in healthy individuals exhibited a lower proportion of IgM+ compared to both cancer and infection groups. This lower ratio in healthy controls reflects a higher baseline proportion of class-switched memory B cells expressing non-IgM isotypes (such as IgG and IgA), whereas active disease states spur fresh recruitment or expansion of unswitched, IgM-expressing lineages.

## Discussion

### Evaluation of cell fractionation in AIRR-seq

In this set of experiments, we partitioned B cells into IgM+ vs. IgM-subsets and T cells into CD4+ vs. CD4-subsets, and evaluated methods to reconstruct the original repertoires for better compatibility with inter-study datasets. Through selective partitioning, we achieved higher resolution and discovered low-abundance clonotypes often overlooked in bulk sequencing assays. We hypothesize that the higher granularity through differential cell sorting adds a layer of analytical depth in diseases which affect a certain subset of T or B lymphocytes, especially in cases where there are larger imbalances in surface marker profiles.

However, selective enrichment creates a methodological barrier when attempting to cross-compare or pool these enriched profiles with legacy, unfractionated datasets. Hence, we investigated the workflow to restore characteristics of the original repertoire from its partitioned subsets for the possibility of integrating partitioned repertoire with external unfractionated datasets. Cell-type ratios and diversity measures from reconstructed repertoires were found to be comparable with original measures, indicating that key repertoire features were conserved.

### Technical challenges and limitations

In our reconstruction, the ratio of T-cells captured through tagged CD4 columns in the LC partition exceeds typical CD4 ratios in healthy individuals. This discrepancy likely results from non-specific binding during anti-CD4 capture. To verify this hypothesis, an alternative method for estimating relative proportions of T-cell markers such as fluorescence imaging is recommended as an orthogonal measure.

The accuracy of the w_FT_ and w_LC_ measurements is an area of concern, particularly given the potential misallocation of lymphocytes intended for the LC partition that may end up in the FT partition due to binding inefficiencies. Our cell sorting method, utilizing magnetic columns, resulted in a 15% to 40% cell loss, averaging around 20%, which impacts the overall yield from the original PBMC. This is further complicated by the use of once-frozen PBMCs, which may have affected the efficiency of both cell sorting and DNA extraction.

When comparing to FACS-sorted cells, isolating for labelled cells as a form of positive selection, leaving the remainder as flowthrough may introduce biases. Assigning CD4+ cells to positive selection while collecting the remainder as flowthrough introduces extraction bias. Incomplete capture or off-target retention systematically skews recovery, yielding higher LC:FT ratios. To mitigate this imbalance, both subsets can be isolated through double-positive immunomagnetic selection. Although MACS recommends a double separation process to enhance purity, this would exacerbate cell loss; given our limited available blood volume, such an approach might compromise DNA yield needed for multiple iPETE replicates.

### Metrics for AIRR-seq analysis

The intrinsic properties of an immune repertoire are often summarized through a series of diversity metrics. In this study, we utilized these indices, specifically Hill numbers, to validate that our reconstructed repertoires maintain the same clonal diversity profiles as the original, unfractionated samples. By applying these metrics, we evaluated the extent to which our reconstruction methods preserve the essential diversity characteristics necessary for downstream analysis. The Bray-Curtis overlap was used to compare reconstructed repertoires, specifically RC1B, against the ground-truth unfractionated TL samples. Higher overlap between the reconstructed repertoire and TL served as an indicator of better reconstruction fidelity.

The ratio of differentially abundant VJ combinations across partitions present an opportunity to identify potential biomarkers for disease states. Notably, infection and cancer subjects exhibit distinct VJ combination abundance profiles compared to healthy individuals, characterized by a shift in the usage of TRB clonotypes expressing CD4 markers as exemplified in Figure S7. While the limited scale of this study does not harness sufficient statistical power to correlate repertoire features with clinical annotations to greater detail, the partition-specific patterns provide valuable insights that may reflect underlying mechanisms of the adaptive immune response.

## Conclusion

This pilot study demonstrated the feasibility of reconstructing adaptive immune receptor repertoires from partitioned T and B cell subsets, providing a valuable method for integrating subset-specific data with broader, unpartitioned AIRRseq datasets. By employing cell-level subsampling streams (RC1B), we successfully preserved key repertoire characteristics, including lymphocyte subtype ratios and clonal diversity, as evidenced by the concordance with original, unfractionated samples. Using RC1B, the method that prioritizes retention of less frequent clones, we addressed technical challenges associated with cell loss during reconstruction.

The identification of differentially abundant VJ combinations, particularly in infection and cancer subjects, suggests potential biomarkers and reflects the dynamic nature of subset-specific immune responses. While this pilot study’s limited scale prevented detailed clinical correlations, the findings underscore the importance of partitioning immune cell subsets for a higher resolution analysis of the immune repertoire. Future studies with larger cohorts and optimized methodologies will further validate these findings and explore the clinical utility of partition-specific repertoire analysis.

## References

Akbari, Vahid, Alexandra Morgan, Michie Yasuda, et al. 2025. “HuBIE: The Human Blood Immunome Encyclopedia of TCRs and BCRs in Bloodstream Infections and Cancer.” In bioRxivorg. October 23. 10.1101/2025.10.22.678684.

Akoglu, Haldun. 2018. “User’s Guide to Correlation Coefficients.” Turkish Journal of Emergency Medicine 18 (3): 91–93.

Carsetti, Rita, Sara Terreri, Maria Giulia Conti, et al. 2022. “Comprehensive Phenotyping of Human Peripheral Blood B Lymphocytes in Healthy Conditions.” Cytometry. Part A : The Journal of the International Society for Analytical Cytology 101 (2): 131–139.

Corrie, Brian D., Nishanth Marthandan, Bojan Zimonja, et al. 2018. “iReceptor: A Platform for Querying and Analyzing antibody/B-Cell and T-Cell Receptor Repertoire Data across Federated Repositories.” Immunological Reviews 284 (1): 24–41.

Donahue, William, Vigneault, and Francois. 2019. Targeted sequencing and UID filtering. US Patent & Trademark Office Patent 10731212, filed June 17, 2019, and issued November 28, 2019.

Gelibter, Alain, Lucrezia Tuosto, Angela Asquino, et al. 2024. “Anti-PD1 Therapies Induce an Early Expansion of Ki67CD8 T Cells in Metastatic Non-Oncogene Addicted NSCLC Patients.” Frontiers in Immunology 15 (December): 1483182.

Gomez-Tourino, Iria, Yogesh Kamra, Roman Baptista, Anna Lorenc, and Mark Peakman. 2017. “T Cell Receptor β-Chains Display Abnormal Shortening and Repertoire Sharing in Type 1 Diabetes.” Nature Communications 8 (1): 1792.

Gros, Alena, Paul F. Robbins, Xin Yao, et al. 2014. “PD-1 Identifies the Patient-Specific CD8+ Tumor-Reactive Repertoire Infiltrating Human Tumors.” The Journal of Clinical Investigation 124 (5): 2246–2259.

Kleiveland, Charlotte R. 2015. “Peripheral Blood Mononuclear Cells.” In The Impact of Food Bioactives on Health: In Vitro and Ex Vivo Models [Internet]. Springer.

Laidlaw, Brian J., Joseph E. Craft, and Susan M. Kaech. 2016. “The Multifaceted Role of CD4(+) T Cells in CD8(+) T Cell Memory.” Nature Reviews. Immunology 16 (2): 102–111.

Lefranc, Marie-Paule, Véronique Giudicelli, Chantal Ginestoux, et al. 2009. “IMGT, the International ImMunoGeneTics Information System.” Nucleic Acids Research 37 (Database issue): D1006–12.

Mark, Michal, Shlomit Reich-Zeliger, Erez Greenstein, et al. 2022. “A Hierarchy of Selection Pressures Determines the Organization of the T Cell Receptor Repertoire.” Frontiers in Immunology 13 (July): 939394.

Mazzotti, Lucia, Anna Gaimari, Sara Bravaccini, et al. 2022. “T-Cell Receptor Repertoire Sequencing and Its Applications: Focus on Infectious Diseases and Cancer.” International Journal of Molecular Sciences 23 (15). 10.3390/ijms23158590.

Miller, Jacqueline Rose, Cynthia Feng, Jordan Ranum, and Rob Striker. 2025. “Viruses Tipping the Scales: The Role of the CD4/CD8 Ratio in Determining Viral Outcome.” Virology 603 (February): 110333.

Olsen, Tobias H., Fergus Boyles, and Charlotte M. Deane. 2022. “Observed Antibody Space: A Diverse Database of Cleaned, Annotated, and Translated Unpaired and Paired Antibody Sequences.” Protein Science 31 (1): 141–146.

Qian, Xinyang, Guang Yang, Fan Li, et al. 2024. “DeepLION2: Deep Multi-Instance Contrastive Learning Framework Enhancing the Prediction of Cancer-Associated T Cell Receptors by Attention Strategy on Motifs.” Frontiers in Immunology 15 (March): 1345586.

Robins, Harlan S., Santosh K. Srivastava, Paulo V. Campregher, et al. 2010. “Overlap and Effective Size of the Human CD8+ T Cell Receptor Repertoire.” Science Translational Medicine 2 (47): 47ra64.

Seo, Kayoung, and Jung Kyoon Choi. 2025. “Comprehensive Analysis of TCR and BCR Repertoires: Insights into Methodologies, Challenges, and Applications.” Genomics & Informatics 23 (1): 6.

