## Supplementary Figures for "Accurate Reconstruction of Adaptive Immune Receptor Repertoires from FACS-partitioned B and T Cell Subsets"

### Extended Figures

**Figure S1** - Schematic workflow of the Daedalus bioinformatics pipeline used to process immune repertoire sequencing data, including steps for quality control, clonotype assembly and gene annotation.

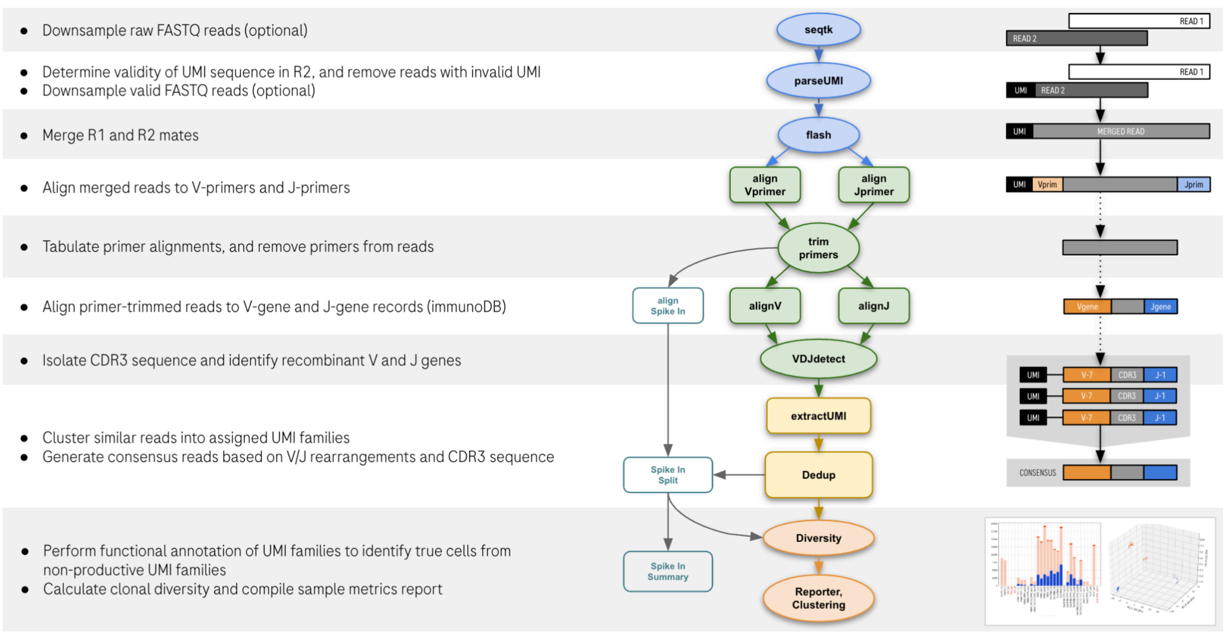


**Figure S2 -** Comparison of CDR3 length distributions in productive versus non-productive families. Most common reasons for non-productive CDR3s were frameshift mutations resulting in a misaligned reading frame (16.5%) and premature stop codons (5%).

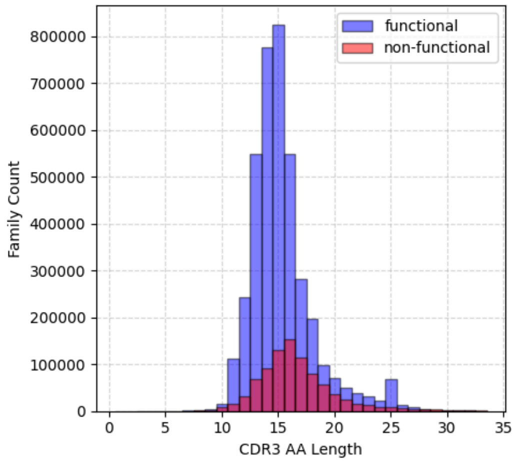


**Figure S3** - Comparison of cell type ratios across three patient groups. The unfractionated TL sample had 81.8% ± 11.7% αβ T-cells, 1.9% ± 2.2% γδ T-cells, and 16.3% ± 12.0% B-cells (IGH chains). The LC fraction showed 93.8% ± 5.3% TRB, while the FT fraction had 62.7% ± 20.4% TRB.

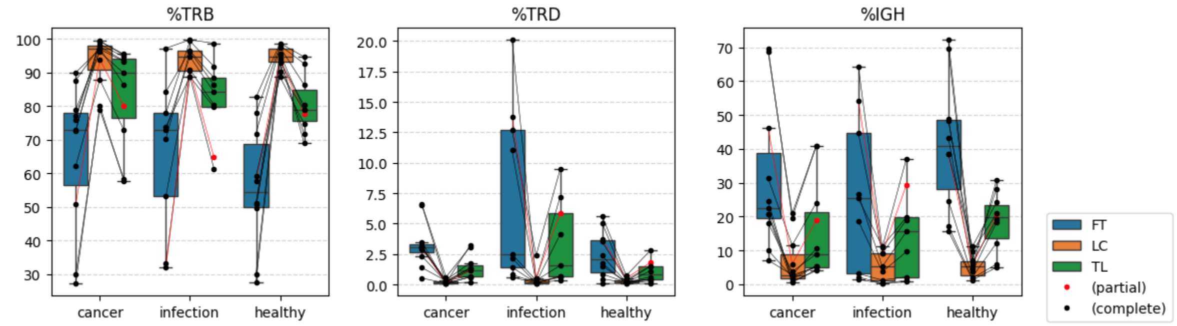


**Figure S4 -** Comparison of Hill numbers through different reconstructions. Reconstruction methods RC1B and RC2A, which retain rare clonotypes, exhibit greater diversity at lower Hill numbers (emphasizing clonal richness). Effective diversity measures align with TL from D2 onwards, where a balance between sample richness and dominant clones is achieved.

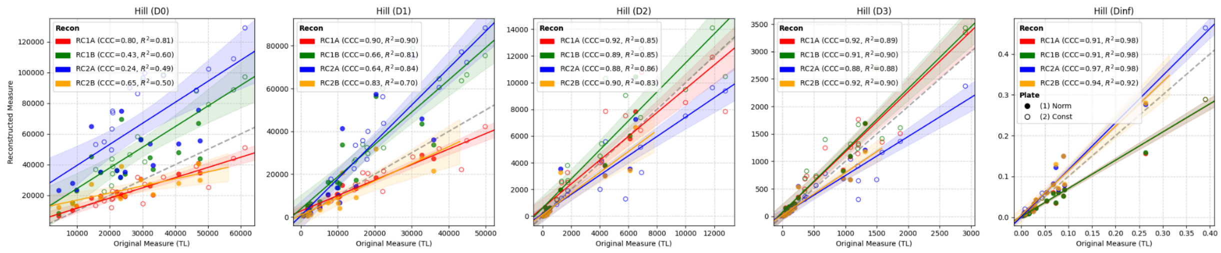

**Figure S5.1 -** The figure compares clonal diversity of reconstructed repertoires (FTLC) through two approaches: (RC1) stratified subsampling at the cell level to match lymphocyte ratios, and (RC2) weighting cell counts based on the LC:FT ratio.

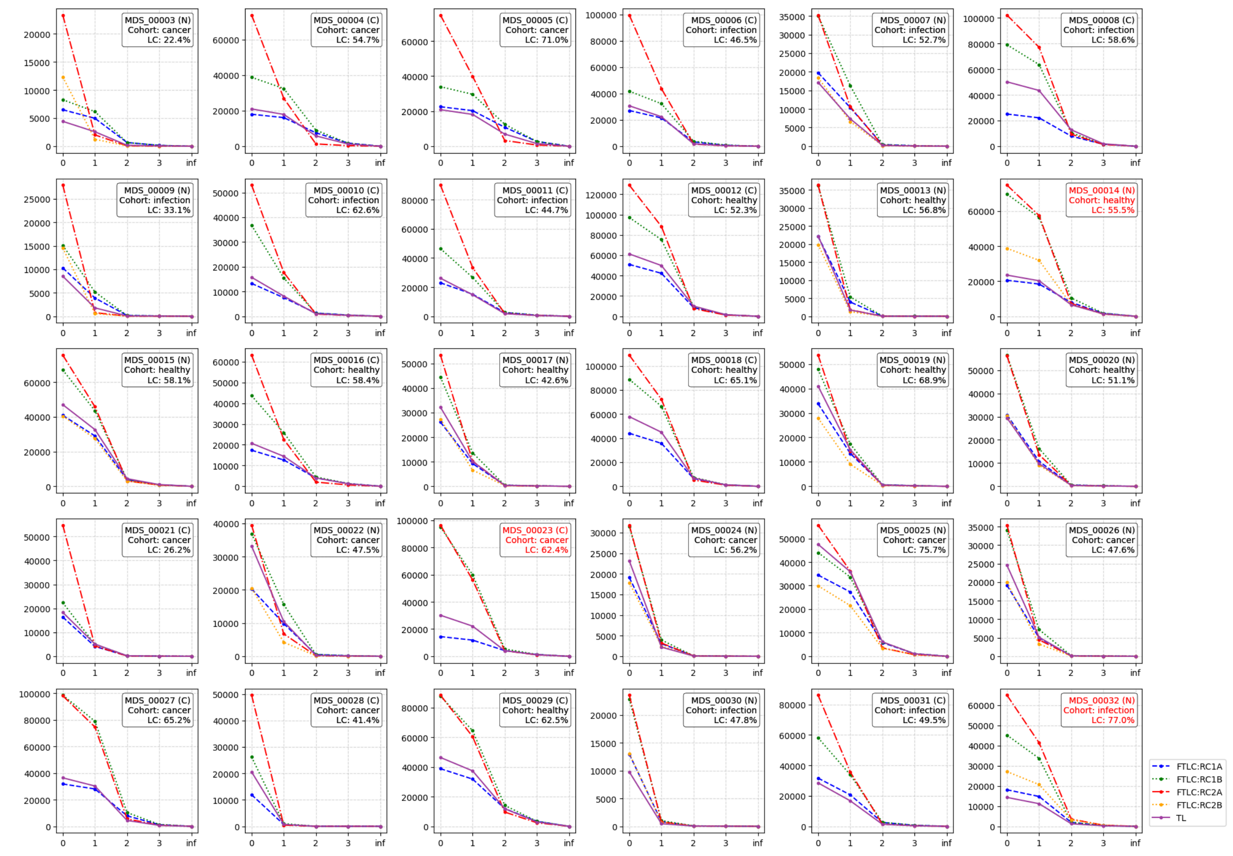


**Figure S5.2 -** The area between the diagonal line and the Lorenz curve, divided by the total area under the diagonal, represents the Gini index

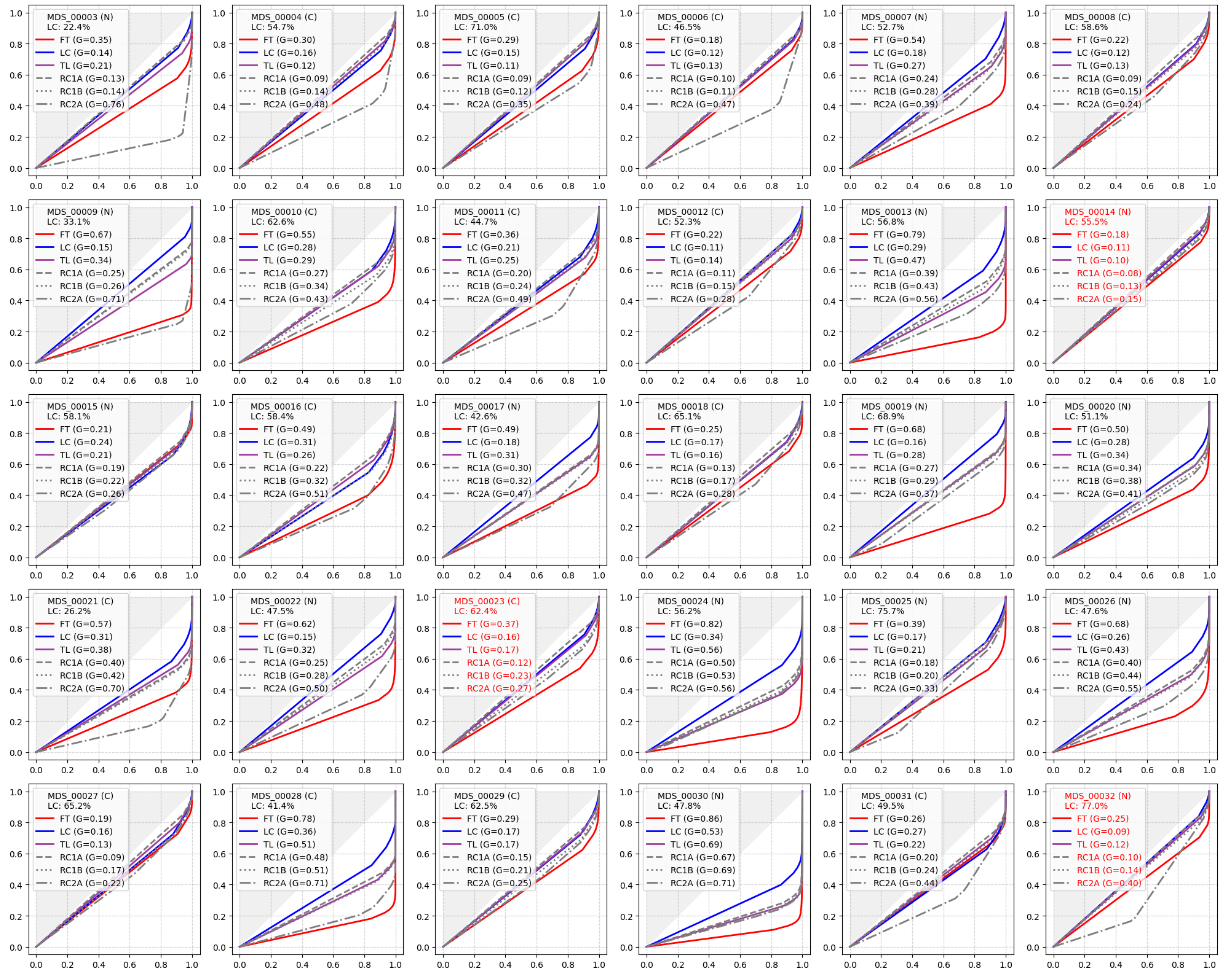


**Figure S6 -** Gene usage plots showing the mean counts per million (cpm)-normalized frequencies of B-cell and T-cell populations for V/J rearrangements, comparing differences between FT and LC partitions.

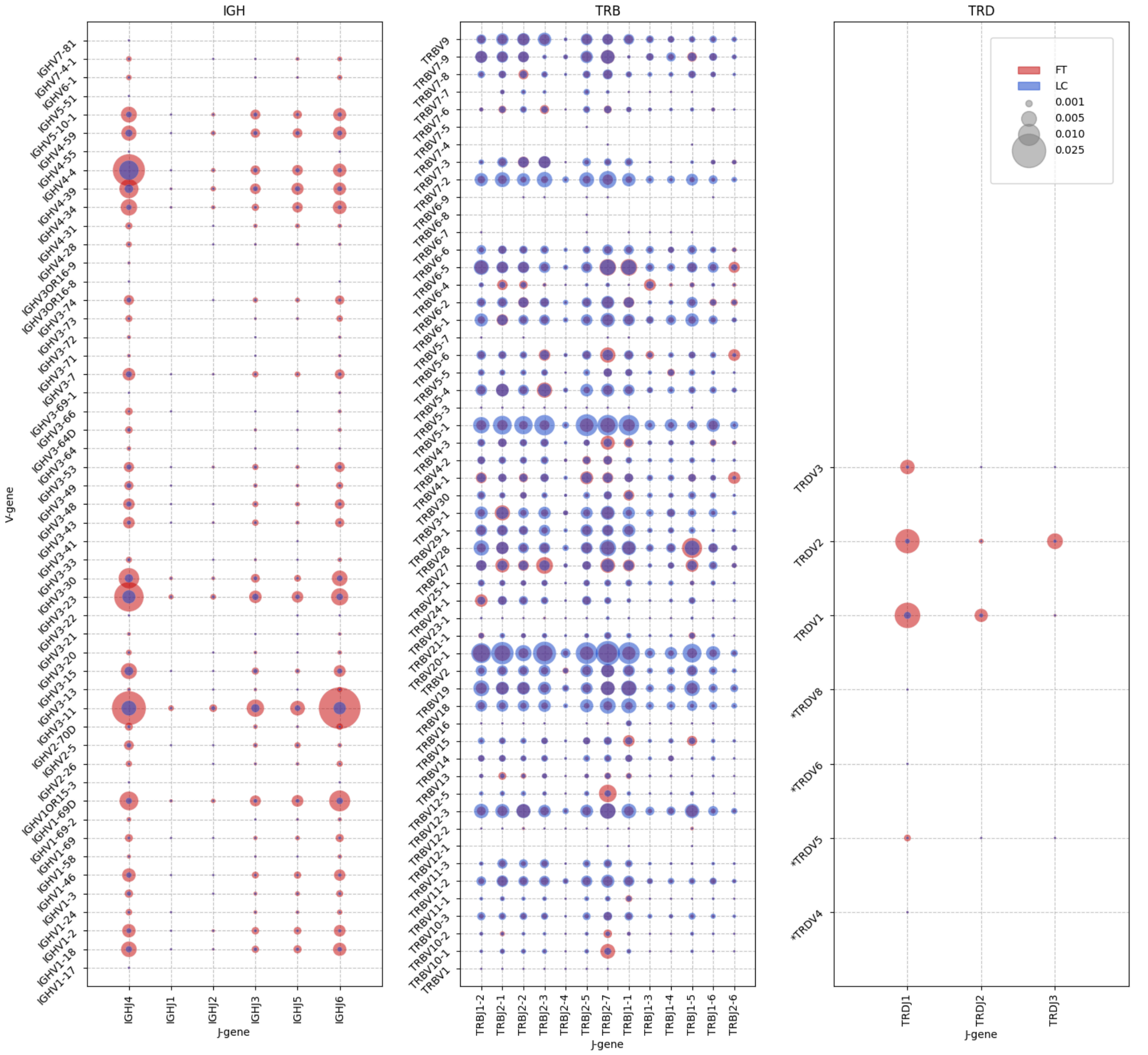


**Figure S7 -** Differential abundance in a larger set of V/J-gene combinations across infection and cancer compared to healthy states observed through a log2 fold-change reflect a potential shift in CD4+ T-cell subpopulations in response to disease-associated antigens.

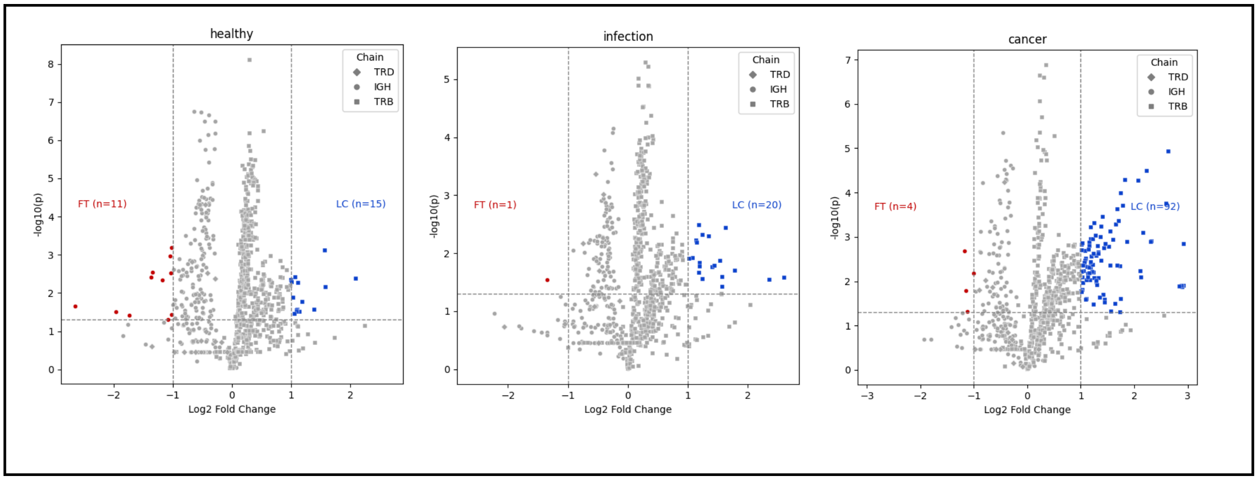

healthy
FT (n=11)
LC (n=15)
-log10(p)
Log2 Fold Change
Chain
TRD
IGH
TRB
infection
FT (n=1)
LC (n=20)
-log10(p)
Log2 Fold Change
Chain
TRD
IGH
TRB
cancer
FT (n=4)
LC (n=92)
-log10(p)
Log2 Fold Change
Chain
TRD
IGH
TRB
